# Small mitochondrial protein NERCLIN regulates cardiolipin homeostasis and mitochondrial ultrastructure

**DOI:** 10.1101/2021.01.03.424667

**Authors:** Svetlana Konovalova, Rubén Torregrosa-Muñumer, Pooja Manjunath, Sundar Baral, Xiaonan Liu, Minna Holopainen, Jouni Kvist, Jayasimman Rajendran, Yang Yang, Markku Varjosalo, Reijo Käkelä, Pentti Somerharju, Henna Tyynismaa

## Abstract

Cardiolipin (CL) is an essential phospholipid for mitochondrial structure and function. Here we present a small mitochondrial protein, NERCLIN, as a negative regulator of CL homeostasis and mitochondrial ultrastructure. Primate-specific NERCLIN is expressed ubiquitously from *GRPEL2* locus on a tightly regulated low level, but induced by heat stress. NERCLIN overexpression severely disrupts mitochondrial cristae structure and induces mitochondrial fragmentation. Proximity labeling suggested interactions of NERCLIN with CL synthesis and prohibitin complexes on the matrix side of the inner mitochondrial membrane. Lipid analysis indicated that NERCLIN regulates mitochondrial CL content. The regulation may occur directly through interaction with PTPMT1, a proximal partner on the CL synthesis pathway, as its product phosphatidylglycerol was also reduced by NERCLIN. We propose that NERCLIN contributes to stress-induced adaptation of mitochondrial dynamics and turnover by regulating the mitochondrial CL content. Our findings add NERCLIN to the group of recently identified small mitochondrial proteins with important regulatory functions.

## INTRODUCTION

Alternative splicing is one of the main sources of protein diversity in higher eukaryotes^1^. Although high throughput techniques routinely detect thousands of alternatively spliced transcripts, proteomics analysis often fails to identify a vast majority of them. Short isoforms of canonical proteins are particularly difficult to detect since the existing algorithms usually exclude them from the proteome annotation. Several recent studies have identified novel small proteins that play important roles in fundamental biological processes such as muscle development and activity^2–5^, mRNA degradation^6^ and transcriptional repression^7, 8^. Interestingly, the mitochondrial proteome is particularly enriched for small proteins^9^, many of which have vital cellular functions. For example, BRAWNIN, a 71-amino acid peptide encoded by *C12orf73*, is essential for vertebrate oxidative phosphorylation^9^, a 54-amino acid mitochondrial microprotein PIGBOS regulates the unfolded protein response^10^, and a 56-amino acid microprotein MTLN (mitoregulin) controls mitochondrial protein complex assembly^11^ and adipocyte metabolism^12^.

Cardiolipin (CL) is a mitochondria-specific phospholipid, which is essential for functional shaping of the mitochondrial inner membrane thus contributing to cristae structure and stabilization of respiratory chain complexes and other components important for bioenergetics^13, 14^. CL is synthesized within the inner membrane by the consecutive actions of phosphatidylglycerophosphate synthase (PGS1), phosphatidylglycerophosphate phosphatase (PTPMT1), and CL synthase (CLS1), followed by fatty acyl chain remodeling to generate mature CL species^15^. PGS1, PTPMT1, and CLS1 form a CL synthesis complex, which interacts with cristae membrane-organizing proteins such as prohibitins^16^. Abnormal CL content or species composition disrupt mitochondrial function and form, and are linked to mitophagy, the selective mitochondrial degradation pathway^14, 15, 17, 18^.

Here we describe the identification of NERCLIN, a human mitochondrial protein of 90 amino acids in its mature form, which is produced as an alternative splice variant of *GRPEL2* gene. GRPEL2 is a nucleotide exchange factor for mitochondrial protein import chaperone mtHSP70^19, 20^, however, NERCLIN does not share the amino acids required for the canonical function of GRPEL2. Instead, we suggest that NERCLIN is a small alpha-helical protein functioning as a negative regulator of CL metabolism and mitochondrial integrity.

## RESULTS

### Identification of NERCLIN, a novel splice variant of human *GRPEL2*

While exploring the human *GRPEL2* gene for our previous study^20^ on the UCSC Genome Browser^21^, we noticed the presence of expressed sequence tags for two separate transcripts. The transcript variant 1 contains four exons, encoding the 225-amino acid GRPEL2 protein, which has a 32-amino acid N-terminal mitochondrial targeting sequence (MTS). The rare transcript variant 2 skips exon 3 producing an exon 2/4 splice junction (Fig. 1a). Absence of exon 3 results in out-of-frame deletion and translational frameshift that potentially generates a 122-amino acid protein, which we termed NERCLIN (negative regulator of CL) (Fig. 1b). Thus, NERCLIN shares the N-terminal MTS and successive 45 amino acids with GRPEL2, but has a unique C-terminus of 45 amino acids, forming a mature protein of 90 amino acids.

**Fig. 1:**
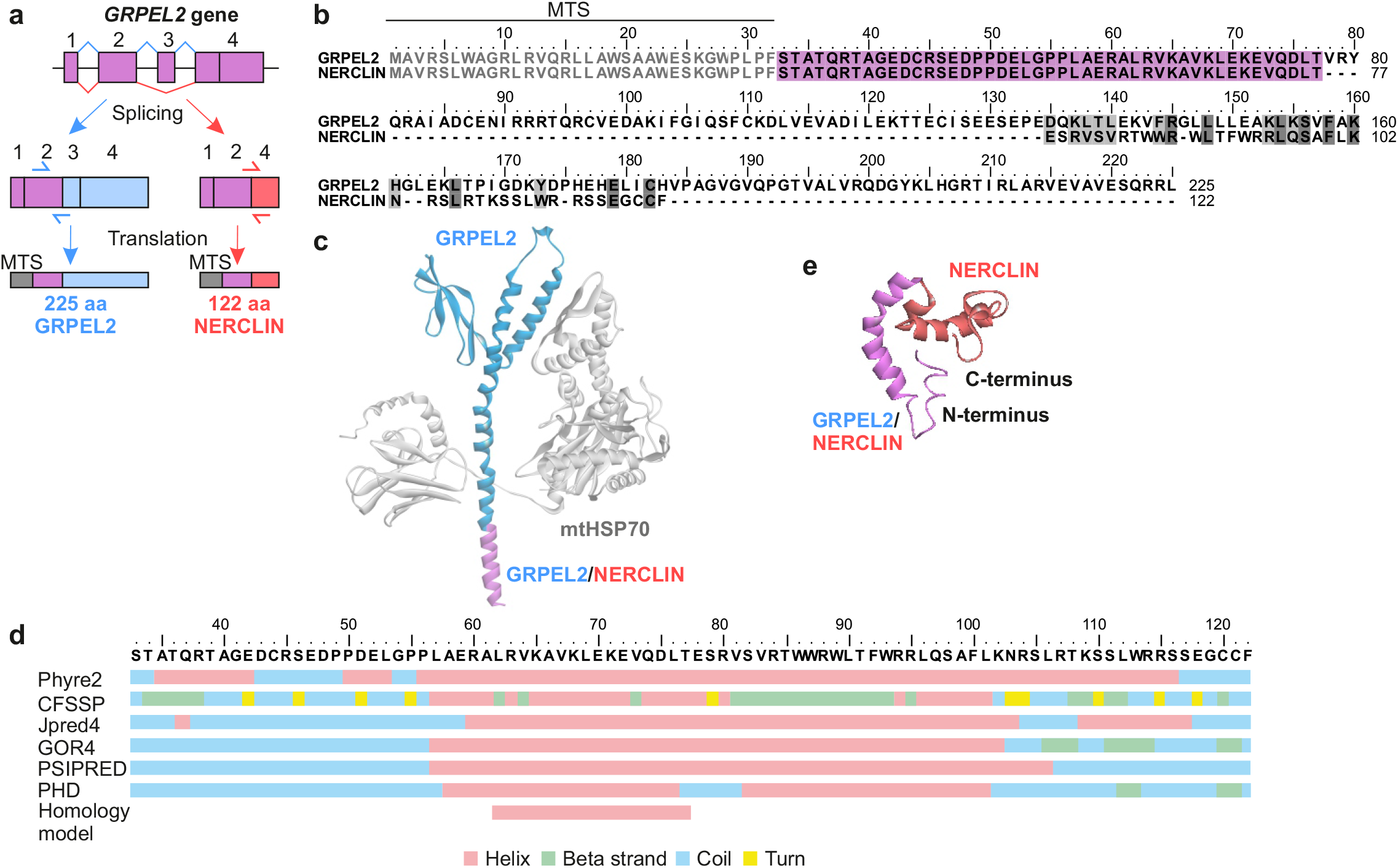
Identification of NERCLIN, a novel splice variant of human *GRPEL2*. **a** Schematic representation of the alternative splicing of the human *GRPEL2* gene, highlighting the parts of the transcripts and proteins that are specific for *GRPEL2* in blue, and for *NERCLIN* in red. The mitochondrial targeting sequence (MTS) is in gray, followed by the part of the protein common for GRPEL2 and NERCLIN in violet. Location of the transcript-specific primer pairs are shown by arrows. **b** Amino acid sequence alignment of human GRPEL2 and NERCLIN. Identical residues are highlighted in dark gray, similar residues are in light gray. Amino acid sequence common for GRPEL2 and NERCLIN is in violet. MTS (1-32 amino acids) is in gray. **c** Structural model of GRPEL2 interacting with mtHSP70 (gray). N-terminal α-helix present in NERCLIN is in violet, the rest of GRPEL2 molecule is in blue. For simplicity, the structure of only one GRPEL2 molecule in GRPEL2 dimer is shown. **d** Secondary structural prediction of human NERCLIN. For the analysis MTS was removed from the protein sequence. **e** Tertiary structure prediction of human NERCLIN by SSpro server. The N-terminal α-helix present in NERCLIN is in violet and the unique amino acid sequence for NERCLIN is in red. MTS was removed from the protein sequence.

We have previously generated a homology model, which shows that the C-terminus of human GRPEL2 interacts with the catalytic ATPase domain of mtHSP70 chaperone^20^. As NERCLIN lacks the critical part for binding mtHSP70, it cannot perform a GrpE-like nucleotide exchange function. Instead, according to the homology model, NERCLIN corresponds to the N-terminal alpha helix of GRPEL2 (Fig. 1c). Structure predictions by several algorithms support that mature NERCLIN is mainly a small alpha helix (Fig. 1d, e). In addition, using Phobius, TMHMM, TMFoldWeb, DAS algorithms, we predicted that NERCLIN does not contain transmembrane domains.

To verify that the *NERCLIN* transcript is expressed in human cells, we performed RT-PCR amplification of RNA from 143B osteosarcoma cells using primers located in *GRPEL2* exons 1 and 4. We observed two amplification products corresponding exactly to the expected amplicons from *GRPEL2* and from *NERCLIN* lacking exon 3 (Supplementary Fig. 1a). Nucleotide sequencing of each product confirmed that the larger transcript corresponded to *GRPEL2* while the smaller transcript was an alternatively spliced *NERCLIN* that contained an out-of-frame deletion of the entire exon 3 comprising 82 bases (nt 232-314) (Supplementary Fig. 1a, b).

Quantitative RT-PCR analysis using primers specific for *GRPEL2* or *NERCLIN* showed that *NERCLIN* was expressed in all tested human tissues (Fig. 2a, b), 13 different human brain regions (Fig. 2c), and in cultured human cells (Fig. 2b), suggesting that it is ubiquitously expressed. We estimate that *NERCLIN* expression is between 1% to 20% of *GRPEL2* expression level, depending on the tissue type or cell line (Fig. 2b, c). The highest relative expression level of about 20% we observed in the placenta, and of the brain regions the highest expression was in the thalamus. To investigate *NERCLIN* expression at stress conditions we subjected HEK293 cells to heat or oxidative stress. We observed a significant induction of *NERCLIN* (up to 2.3 fold change) in heat stress (Fig 2D), but not in oxidative stress (Supplementary Fig. 1c), while *GRPEL2* expression was unchanged. Thus *NERCLIN* expression is responsive to heat stress.

**Fig. 2:**
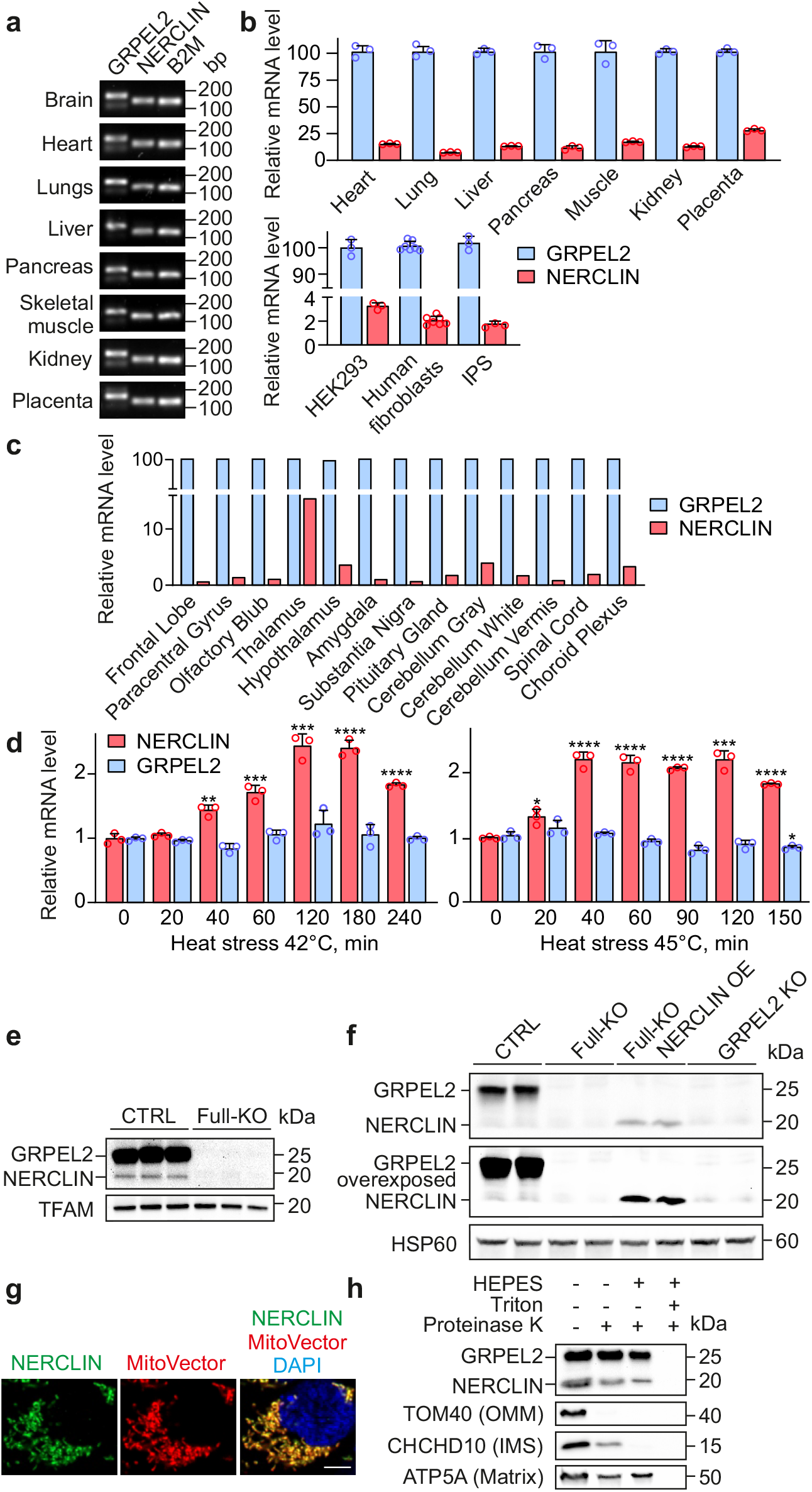
NERCLIN is expressed in human cells. **a** Expression pattern of *NERCLIN* mRNA in human tissues. Transcript-specific primer pairs indicated in **Fig. 1a** were used for PCR reaction. Gel electrophoresis of real-time PCR products. β2 microglobulin (B2M) was used as a loading control. **b** Relative mRNA levels of *NERCLIN* in human tissues and cultured cells as determined by qPCR. mRNA level of *GRPEL2* was taken as 100%. n = 3 (human tissues, HEK293 cells or human fibroblasts), n = 5 (IPS). **c** Relative mRNA levels of *NERCLIN* in human brain samples as determined by qPCR. mRNA level of *GRPEL2* was taken as 100%. n = 1. **d** mRNA levels of *NERCLIN* or *GRPEL2* in HEK293 cells exposed to heat stress (42°C or 45°C) as determined by qPCR assay (n = 3). Data are presented as mean ± SD. **P < 0.01, ***P < 0.001, ****P < 0.0001, ns, not significant as compared to control, untreated cells (unpaired t-tests). **e, f** Protein expression of NERCLIN in mitochondria isolated from HEK293 cells. Full-KO cells lacking GRPEL2 and NERCLIN were used as negative control. TFAM or HSP60 was used as a loading control. **g** Intracellular localization of NERCLIN in 143B cells as shown by fluorescent microscopy. NERCLIN-EGFP construct and MitoVector were transiently co-expressed for 24 h. MitoVector was used for labeling mitochondria, DAPI shows nuclei. Scale bar, 5 μm. **h** Mitochondria isolated from HEK293 cells overexpressing NERCLIN were treated with proteinase K at isoosmotic (without HEPES) or hypoosmotic (with HEPES) conditions. OMM, outer mitochondrial membrane, IMS, intermembrane space.

*GRPEL2* gene is found in vertebrates, however, EST databases contained transcripts lacking exon 3 only in some primates. We performed *in silico* analysis with genomic sequences to identify the nonhuman primates that had the potential for a conserved NERCLIN C-terminus following exclusion of exon 3. We found sequences for a highly conserved C-terminus in all apes and in some Old World monkeys, and a less conserved C-terminus in some New World monkeys, tarsiers, and strepsirrhines (Supplementary Fig. 1d, e). As green monkey was among the positive Old World monkeys, we performed RT-PCR analysis of green monkey kidney COS7 cells, and indeed detected the *NERCLIN* transcript lacking exon 3 (Supplementary Fig. 1f). On the contrary, in RNA samples from dog or mouse only the product corresponding to *GRPEL2* could be amplified. These results suggest that NERCLIN is specific to a subgroup of primates.

Next, we investigated whether NERCLIN is effectively translated into a stable protein *in vivo*. We performed western blotting analysis of mitochondria isolated from HEK293 cells using an antibody directed against the GRPEL2 N-terminus (residues 3-115). In addition to the 25 kDa GRPEL2 band, we detected a 15 kDa band putatively corresponding to NERCLIN (Fig. 2d). Importantly, both bands were absent in a full knockout (Full-KO) cell line of *GRPEL2* and *NERCLIN*, which we had previously generated by deleting exon 1 using CRISPR/Cas9^20^. To confirm that the second band was not a result of GRPEL2 protein degradation, we generated a *GRPEL2-specific* knockout cell line (GRPEL2-KO) in HEK293 cells by deleting the exon 3 using CRISPR/Cas9 (Supplementary Fig. 2a, b). In GRPEL2-KO cells, GRPEL2 band was absent while the 15 kDa band stayed intact (Fig. 2e). As *NERCLIN* does not have any unique sequence, it is not possible to generate a cell line lacking NERCLIN without affecting GRPEL2. However, transient overexpression of *NERCLIN* in Full-KO cells confirmed that the 15 kDa band in western blot was NERCLIN (Fig. 2f). In line with the transcript levels, endogenous NERCLIN protein is less abundant than GRPEL2.

NERCLIN shares the N-terminal MTS with GRPEL2 suggesting that NERCLIN is also a mitochondrial protein. Using fluorescent microscopy, we showed that NERCLIN-GFP localizes strictly to mitochondria in human cells (Fig. 2g). To determine the sub-mitochondrial localization of NERCLIN we used protease protection assay. NERCLIN was protected from protease after selective opening of the outer mitochondrial membrane suggesting that NERCLIN localizes to the mitochondrial matrix (Fig. 2h).

Altogether these results show that NERCLIN is a small mitochondrial protein, which is expressed ubiquitously from *GRPEL2* locus, but on a relatively low level, and found specifically in humans and some nonhuman primates.

### BioID analysis of proximal proteins suggests distinct functions for GRPEL2 and NERCLIN

To gain insight into the function of NERCLIN in human cells, we analyzed its proximal proteins by the proximity-dependent biotin identification (BioID) approach^22^. Promiscuous biotin ligase BirA* was fused to the C-terminus of NERCLIN and the expression of the fusion construct in human 143B cells was confirmed by immunoblotting (Fig. 3a). Immunocytochemistry showed that NERCLIN-BirA* fusion protein was localized to mitochondria (Fig. 3b). To exclude non-specifically labeled proteins we used BirA* fused to green fluorescent protein (GFP-BirA*) or to apoptosis inducing factor (AIF-BirA*), a mitochondrial intermembrane space protein, as controls. Immunoblotting analysis following biotin labeling revealed that while the pattern of biotinylated proximal proteins for GRPEL1-BirA* and GRPEL2-BirA* were identical, confirming our previous findings^20^, the pattern for NERCLIN-BirA* was different (Fig. 3c).

**Fig. 3:**
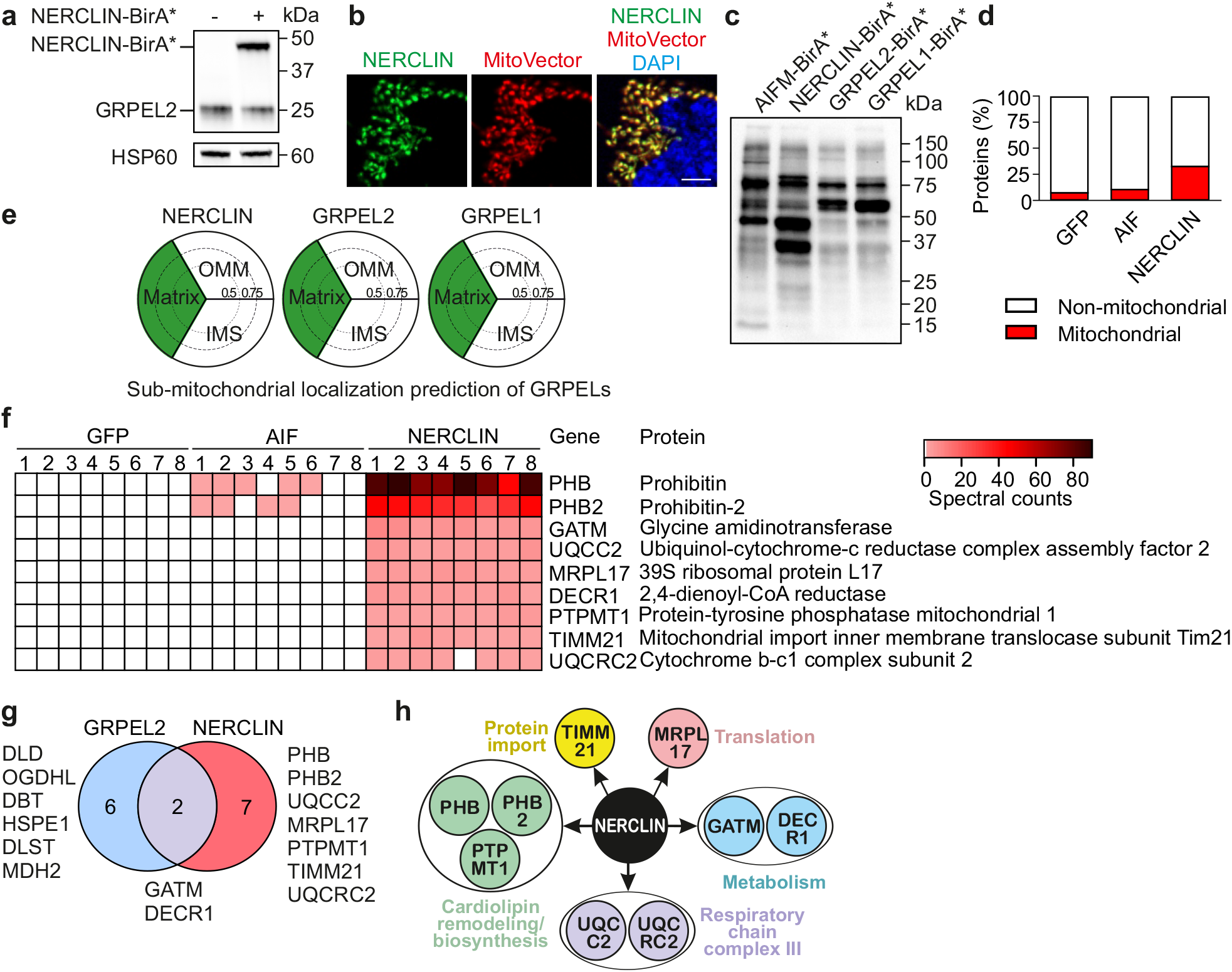
Proteins proximal to NERCLIN identified by BioID. **a** Immunoblotting analysis of NECLIN-BirA* expression in 143B cells. HSP60 was used as a loading control. 143B cells were transiently transfected with indicated constructs for 24 h. **b** Intracellular localization of NERCLIN-BirA* determined by immunocytochemistry. MitoVector was used to label mitochondria, DAPI shows nuclei. Scale bar, 10 μm. **c** Biotinylated proteins in total cell lysates analyzed by immunoblotting. 143B cells were transiently transfected for 24 h with indicated constructs. Biotin was added to the culture media for 24 h. **d** Percentage of the mitochondrial location of all significant proximal proteins of NERCLIN before filtering. **e** Interaction profile identified by BioID analysis was used for MS-microscopy to predict sub-organelle localization of NERCLIN. Three sectors in each plot represent matrix, intermembrane space (IMS) and outer mitochondrial membrane (OMM). Sector areas indicate the possible location score of the proteins, with scores between 0 and 0.5, between 0.5 and 0.75, and 0.75 and 1. (see ^23^ for the reference). **f** Heat map of the unique proximal proteins of NERCLIN. **g** Venn diagram showing distinct unique proximal proteins for GRPEL2 and NERCLIN. **h** Classification of the proteins potentially interacting with NERCLIN.

Biotinylated proteins from four replicates for each BirA* fusion protein were extracted using streptavidin beads and analyzed by mass spectrometry (Supplementary Table 1). Two or more spectral matches in at least three replicates were considered as significant hits and included in further analysis. NERCLIN-BirA* showed enrichment in proximal proteins localized to mitochondria (33%) compared to GFP-BirA* (10%) or AIF-BirA* (12%) (Fig. 3d). Using the BioID data and a previously described algorithm for mapping of protein localization^23, 24^, we confirmed that similarly to GRPEL1 and GRPEL2, NERCLIN localizes to the mitochondrial matrix (Fig. 3e).

To identify highly specific proximal proteins of NERCLIN, we first excluded proteins detected in GFP or AIF control samples. Next, we excluded the mitochondrial matrix BioID matches that are commonly found with any matrix bait^20, 23^. The resulting list of unique highly specific hits consisted of nine proteins that potentially interact with NERCLIN (Fig. 3f). NERCLIN did not show proximity to GRPEL2 since no peptide-spectrum matches specific for GRPEL2 were identified in NERCLIN-BirA* samples (Supplementary Fig. 3). This suggests that GRPEL2 and NERCLIN do not cross interact, although GRPEL2 is known to form homodimers^20^.

Next, we compared the proximal proteins of GRPEL2 and NERCLIN. We used BioID data for GRPEL2 from our previous study where the same filtering was applied^20^. Only two proteins were identified as shared potential interactors for GRPEL2 and NERCLIN, whereas six proteins were specific only to GRPEL2 and seven to NERCLIN (Fig. 3g). These findings suggested that GRPEL2 and NERCLIN function in distinct pathways in human mitochondrial matrix.

Pathway analysis of the potential proteins interacting with NERCLIN revealed that three out of nine proteins were associated with CL synthesis or organization including both prohibitins (PHB and PHB2) and the mitochondrial protein-tyrosine phosphatase (PTPMT1) (Fig. 3f, h). NERCLIN also showed proximity to some respiratory chain complex III and mitochondrial protein import and translation proteins.

### Overexpression of NERCLIN disrupts mitochondrial morphology

BioID data showed high spectral counts for the proximity of NERCLIN to PHB and PHB2. These two prohibitins assemble into heterodimers that form large ring-like complexes at the inner mitochondrial membrane, providing a scaffold for proteins and lipids that control cristae morphogenesis and functional integrity of mitochondria^25–27^. We hypothesized that NERCLIN may affect mitochondrial morphology by its association with the prohibitins. We thus used electron microscopy on Full-KO cells, and on cells overexpressing *NERCLIN*. Interestingly, we observed that while cells without NERCLIN had a comparable mitochondrial morphology to non-transfected cells or cells transfected with empty vector (pBabe), the cells overexpressing *NERCLIN* had a severe disruption of mitochondrial ultrastructure (Fig. 4a). Particularly in *NERCLIN* overexpressing cells, the mitochondria were small and fragmented, and had disordered cristae and irregular intermembrane and intercristal space (Fig. 4a, b). Depletion of prohibitins is known to cause such a phenotype in cultured cells^28, 29^. Next, we tested by immunoblotting if PHB or PHB2 levels were affected in cells overexpressing *NERCLIN*. The levels of all tested mitochondrial proteins were somewhat reduced after 48 h of overexpression, indicating that mitochondrial mass was decreasing in *NERCLIN* expressing cells (Fig. 4c, d). Nevertheless, the immunoblotting results did not indicate that NERCLIN would primarily destabilize prohibitins, as we did not detect a decrease in prohibitin levels relative to the mitochondrial marker TOM40 (Fig. 4c, d) or at equal loading of proteins from isolated mitochondria (Fig. 4c, f). Consistent with the morphological changes induced by NERCLIN, immunoblotting analysis showed that *NERCLIN* overexpression reduced the level of long OPA1 (Fig. 4c, e). In addition, we observed decreased levels of OXPHOS subunits in mitochondria isolated from cells overexpressing *NERCLIN* (Fig. 4c, f). Autophagy marker p62 was unchanged, whereas LC3B-II was reduced by *NERCLIN* overexpression. (Fig. 4c, d).

**Fig.4:**
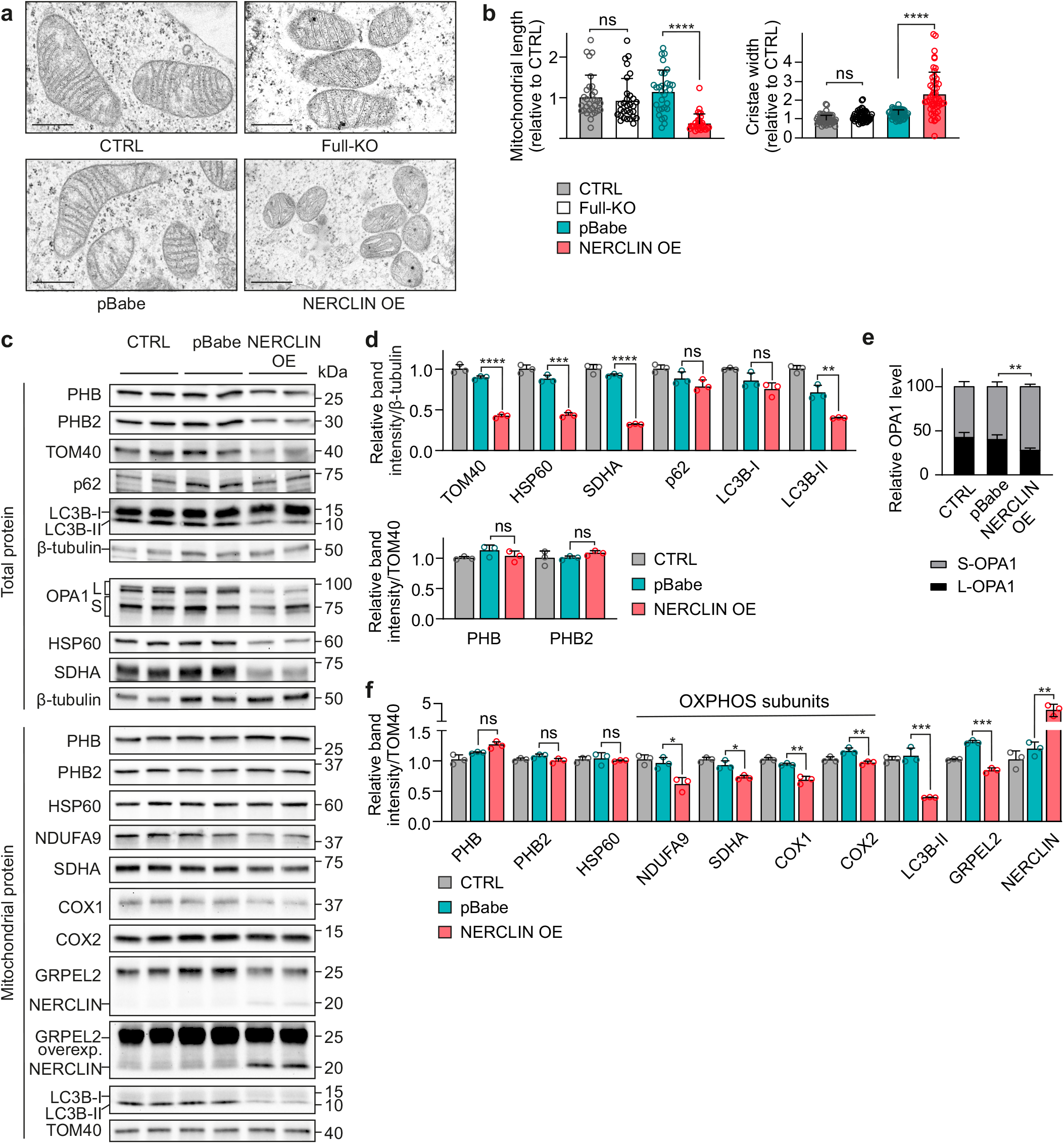
Overexpression of NERCLIN disrupts mitochondrial morphology and ultrastructure. HEK293 cells were transiently transfected with NERCLIN plasmid (NERCLIN OE) or with an empty vector (pBabe) for 48 h. CTRL, non-transfected cells. **a** Mitochondrial ultrastructure in HEK293 cells analyzed by transmission electron microscopy. Full-KO, cells lacking GRPEL2 and NERCLIN. Scale bar: 500 nm. **b** Quantitative analyses of mitochondrial major axis length (mitochondrial length, n = 30) and cristae width (n = 50) from electron microscopy images. **c** Western blot analysis of cells overexpressing NERCLIN. Total protein lysates or mitochondrial protein lysates were used. L, long OPA1 isoform, S, short OPA1 isoform. **d** Quantification of western blot images presented in **(c)**. Protein expression levels are normalized to β-tubulin level (n = 3). **e** Relative levels of short and long OPA1 isoforms determined by western blot analysis in **(c)**. L, long OPA1 isoform, S, short OPA1 isoform (n = 4). **f** Quantification of western blot images presented in **(c)**. Protein expression levels are normalized to TOM40 level (n = 3). Data are shown as mean ± SD. *P < 0.05, **P < 0.01, ***P < 0.001, ****P < 0.0001, ns, not significant as compared to the control cells or cells transfected with empty vector (one-way ANOVA in **b**; unpaired t-tests in **d-f**).

Full-KO cells or the KO cells stably overexpressing *GRPEL2* did not show changes in mitochondrial mass or in the levels of OXPHOS subunits (Supplementary Fig. 2c, d), indicating that in contrast to *NERCLIN* overexpression, which causes severe disruption of mitochondrial ultrastructure, the lack of NERCLIN does not cause mitochondrial abnormalities.

### Overexpression of NERCLIN reduces CL

Prohibitins interact with the CL synthesis complex of which we also detected phosphatase PTPMT1 as a proximal protein of NERCLIN. Deficiency of PTPMT1 has been previously shown to compromise mitochondrial respiration and morphology by depleting phosphatidylglycerol (PG), which is the final substrate for the biosynthesis of CL^30^. Next, we performed lipidomics analysis on cells overexpressing *NERCLIN* to investigate if CL maintenance was affected. Principal component analysis of lipid profiles showed that transfection (empty vector or NERCLIN) had a major effect on cellular lipids. However, cells overexpressing *NERCLIN* also separated from the cells transfected with empty vector (pBabe) based on their lipid profile (Fig. 5a). Interestingly, the analysis of 189 individual lipid species from 11 lipid classes showed a significant reduction of six CL species in cells overexpressing *NERCLIN* compared to the cells transfected with empty vector (Fig. 5b, c), while none of the other lipid species were significantly altered. Analysis of the CL profile suggested no accumulation or depletion of a specific CL species upon overexpression of *NERCLIN* (Fig. 5d), but a general reduction in CL content. This observation suggested that the overall CL synthesis was affected by NERCLIN, and not a particular step in CL maturation. Since CL is located almost exclusively in the mitochondrial inner membrane, the reduced cellular CL level could reflect the decrease of mitochondrial content, which we had observed by western blotting of mitochondrial proteins in cells overexpressing NERCLIN (Fig. 4c, e). To exclude that the reduction in CL level was solely an indication of the decreased mitochondrial mass, we measured total CL in HEK293 cells and in isolated mitochondria using a fluorometric assay. We found CL level to be reduced both in total cells and in isolated mitochondria as a result of *NERCLIN* overexpression (Fig. 5e), indicating that NERCLIN specifically depleted mitochondrial CL. Overexpression of *NERCLIN* for 48 h was required to detect the changes in CL content or in OPA1 isoforms (Supplementary Fig. 4 a-d). In line with the unaltered mitochondrial morphology in the absence of NERCLIN, the levels of CL species were normal in Full-KO cells (Supplementary Fig. 2 e). *NERCLIN* overexpression also fragmented mitochondria in human fibroblasts and in green monkey COS7 cells, as determined by OPA1 western blot (Supplementary Fig. 5 a,b,d,e,g). In addition, CL content was decreased in human fibroblasts overexpressing *NERCLIN* (Supplementary Fig. 5h). In contrast, mouse embryonic fibroblasts overexpressing *NERCLIN* had no changes in CL content or in the levels of OPA1 isoforms (Supplementary Fig. 5 a,c,f,h). These data indicate that NERCLIN effect on mitochondria and cardiolipin is primate-specific.

**Fig. 5:**
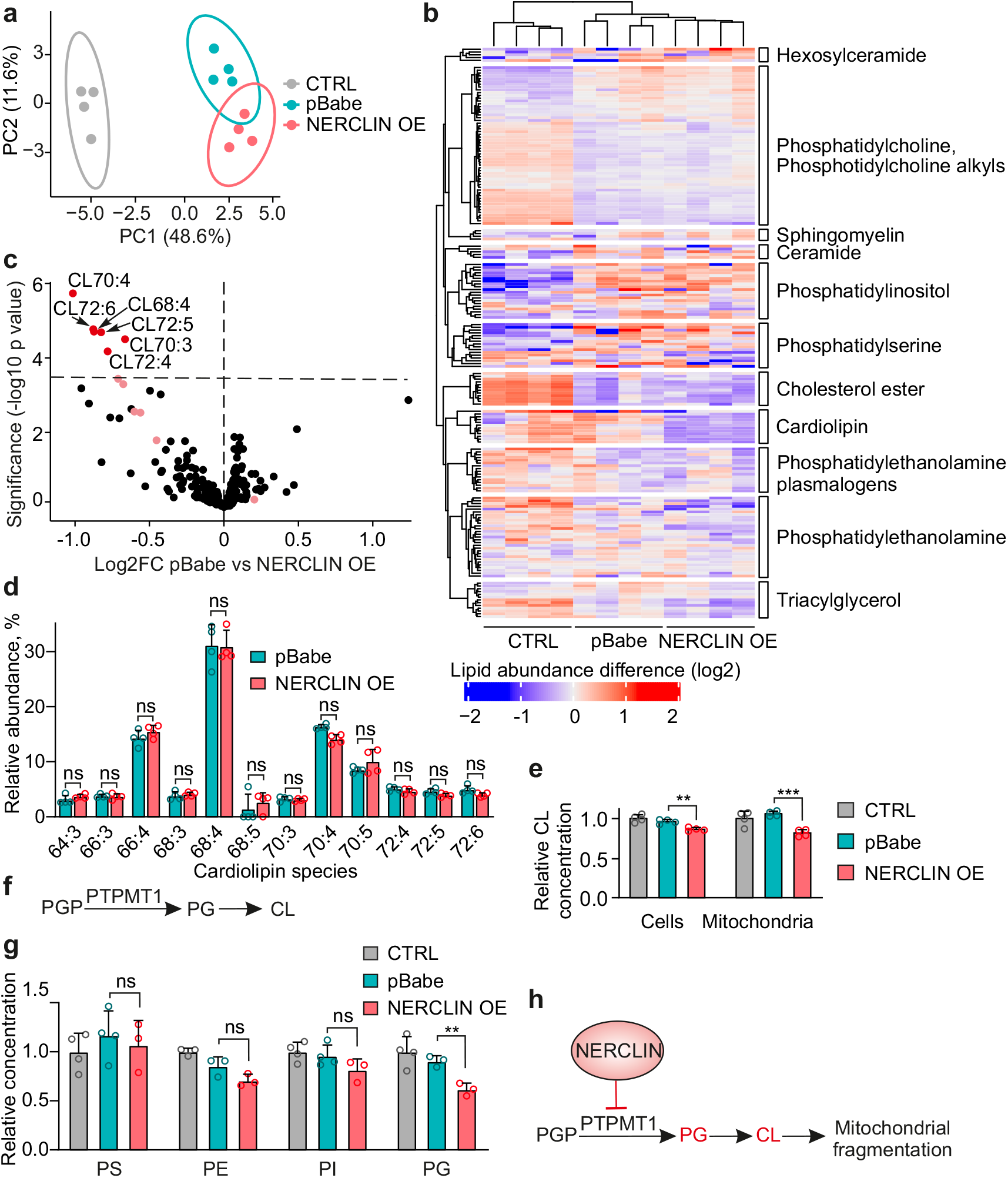
Overexpression of NERCLIN reduces CL level. HEK293 cells were transiently transfected with NERCLIN plasmid (NERCLIN OE) or with an empty vector (pBabe) for 48 h. Lipids from four biological replicates for each sample were extracted and analyzed by mass spectrometry. CTRL, non-transfected cells. **a** Principal component analysis (PCA) of control (CTRL), NERCLIN overexpressing cells and cells transfected with empty vector (pBabe). **b** Heat map showing the relative lipid concentrations of individual species in control (CTRL), NERCLIN overexpressing cells and cells transfected with empty vector (pBabe). Each column represents one sample and each line represents one compound colored by its normalized abundance. Lipid abundance ratios are colored according to the fold changes and the color key indicates the magnitude of log2 fold change. **c** Volcano plot compares the lipid composition of NERCLIN overexpressing cells and cells transfected with empty vector (pBabe). The dashed line indicates a false discovery rate (FDR)-corrected P-value of 0.05. Significantly changed lipids are in red, others in black. Light red indicates CL species that are not significantly changed. **d** CL profile of NERCLIN overexpressing cells compared to cells transfected with empty vector (pBabe). The abundance of individual CL species is normalized to total CL content. **e** Total CL concentration in HEK293 cells or in mitochondria isolated from HEK293 cells transfected with NERCLIN or empty vector (pBabe) as determined by fluorometric assay (n = 4). **f** Scheme showing PTPTMT1 role in CL biosynthesis pathway. **g** Phospholipid class relative concentrations measured by mass spectrometry in mitochondrial fractions and normalized against CTRL values. PS, phosphatidylserine; PE, phosphatidylethanolamine; PI, phosphatidylinositol; PG, phosphatidylglycerol (n = 3–4). **h** Putative mechanism of NERCLIN in the regulation of mitochondrial morphology. We propose that NERCLIN is a negative regulator of CL biosynthesis. By inhibiting PTPMT1 activity, NERCLIN causes depletion of phosphatidylglycerol, an essential precursor of CL. The decrease of CL level destabilizes mitochondrial morphology and induces mitochondrial fragmentation that in turn can have a protective role under stress conditions. PGP, phosphatidylglycerol phosphate, PG, phosphatidylglycerol. In all graphs data are presented as mean ± SD. **P < 0.01, ***P < 0.001, ns, not significant as compared to the cells transfected with empty vector (unpaired t-tests).

As PTPMT1 catalyzes the dephosphorylation of PG-phosphate (PGP) to generate PG (Fig. 5f)^30, 31^, we measured PG levels in mitochondria isolated from cells overexpressing *NERCLIN*. Interestingly, PG level was reduced in mitochondria of *NERCLIN* overexpressing cells, whereas phosphatidylethanolamine (PE), phosphatidylinositol (PI), or phosphatidylserine (PS) levels were not changed (Fig. 5g). Thus, the lipid analysis suggest that NERCLIN negatively regulates CL biosynthesis by inhibiting PTPMT1 activity.

## DISCUSSION

We have here identified and characterized a previously unknown human mitochondrial matrix protein, NERCLIN. It represents a new small protein, being 90 amino acids in its mature form after MTS cleavage. NERCLIN is transcribed from the *GRPEL2* locus, however, it does not have the properties of a GrpE-like nucleotide exchange factor. Instead, NERCLIN is likely to be a small alpha-helical protein, which we find to locate in the proximity of the prohibitins and the CL synthesis complex at the matrix side of the inner mitochondrial membrane. NERCLIN is ubiquitously expressed in human cells and tissues, and its normally relatively low expression level appears to be tightly regulated. However, we find that *NERCLIN* transcription is induced by heat stress, suggesting that it has a role in stress-induced regulation of mitochondrial structure and function. The absence of NERCLIN does not cause any obvious defects in mitochondria of cultured cells, but its overexpression leads to dramatic changes in mitochondrial structure and cristae maintenance. These changes associate with a specific reduction in CL levels. Thus we propose NERCLIN as a negative regulator of CL and mitochondrial ultrastructure.

Our unbiased BioID and lipidomics results point to NERCLIN having major effects on mitochondria through its interactions in the site where prohibitin complexes form scaffolds for CL synthesis^32^. Another recently identified small mitochondrial protein, mitoregulin, is also linked to CL. It is a single pass transmembrane protein rich in positively charged arginine and lysine residues, which CL binds^11^. In contrast to NERCLIN, mitoregulin has positive effects on mitochondrial functions, such as on respiratory supercomplex assembly, and it is found widely in vertebrates. NERCLIN has also several arginines and lysines, which may enable direct binding to CL. However, the lower level of PTPMT1 product PG suggested that NERCLIN may directly inhibit its enzyme activity. Loss of PTPMT1 results in mitochondrial fragmentation and abnormal mitochondrial morphology similarly to the phenotype that we observed in cells overexpressing *NERCLIN*^30^. Prohibitin levels were not affected, suggesting that besides being in close proximity to prohibitins, NERCLIN does not interfere with their assembly.

Mitochondrial proteome is enriched in small proteins, which was speculated to be energetically favorable considering the need to import the proteins through the double membranes^9^. Another reason may be the wide requirement for regulation of mitochondrial form and function depending on a tissue and cell state, for which a range of small proteins may have evolved to contribute to. CL has a profound importance for mitochondrial function and dynamics, requiring a complex regulation, and thus NERCLIN may have evolved to adjust CL biosynthesis to precise cellular needs. Why NERCLIN is specific to primates and what its physiological role is in primate cells and tissues are open questions. Interestingly, *TAZ* encoding mitochondrial acyltransferase taffazin, which mediates CL maturation, also has a primate-specific splice variant^33, 34^. The induction of *NERCLIN* expression by heat may provide a clue for its specific function. In *Arabidopsis*, CL was shown to mediate mitochondrial dynamics and heat stress response^35, 36^. We propose that stress-induced elevation of NERCLIN inhibits CL biosynthesis and thus causes mitochondrial fragmentation (Fig. 5h). Mitochondrial fission and fragmentation have an adaptative and protective role in stress response maintaining bioenergetics homeostasis^37, 38^, but in the excess may promote mitochondrial degradation. CL has also been shown to directly serve as an elimination signal for mitophagy when it is externalized to the outer mitochondrial membrane, where it interacts with LC3B^39^. In NERCLIN overexpressing cells, mitochondrial mass as well as lipidated LC3B-II levels were decreased, which may suggest active turnover of mitochondria. Further studies are needed to elucidate the precise function of NERCLIN, however, our results indicate why its expression must be strictly controlled.

In conclusion, NERCLIN identified in this study significantly differs from GRPEL2 by its expression, structure, function, and regulation in stress conditions. Similarly to many other small proteins, NERCLIN has been overlooked in the previous transcriptomic and proteomic studies due to its small size and partial overlap with GRPEL2 gene and protein.

## MATERIALS AND METHODS

### Cell culture

Human embryonic kidney cells (HEK293), human osteosarcoma cells (143B), human fibroblasts, mouse embryonic fibroblasts (MEFs) or African green monkey kidney fibroblasts (COS7) were cultured (37°C, 5% CO2) in DMEM (Lonza), supplemented with 10% fetal bovine serum (Life Technologies), l-glutamine (Gibco) and penicillin/streptomycin (Gibco). For heat shock treatments, cells were incubated at 45°C or 42°C for indicated times and collected immediately after heat shock. JetPrime reagent (Polyplus) was used for cell transfection. After transfection cells were incubated for 24 or 48 h. To generate stably overexpressing cell lines retroviral constructs were transiently transfected into phoenix package cell line with jetPRIME transfection reagent (Polyplus) and HEK cells were infected by being exposed to virus-containing medium in the presence of polybrene. Infected cells were treated with 1.5 μg/ml puromycin (Sigma) for five days until resistant clones formed. For heat shock treatments, cells were kept in the incubator at 45°C for the indicated time and collected immediately after heat shock. To induce oxidative stress cells were treated with 250 μM or 500 μM H_2_O_2_ (Sigma) for the indicated time.

### Mitochondria isolation

Cultured HEK293 cells were harvested at 1000 g for 10 min at 4 °C. Cell pellets were resuspended in ice-cold mitochondria isolation buffer (200 mM mannitol, 70 mM sucrose, 10 mM HEPES, 1mM EGTA, 0.2 % delipidated BSA, protease inhibitor cocktail (Thermo Scientific), pH 7.5) and homogenized with 60 strokes using Dounce glass-glass homogenizer. Nuclei and unbroken cells were removed by centrifugation at 600 g for 20 min. The supernatant was centrifuged at 10 000 g for 10 min to pellet mitochondria. Mitochondrial pellets were resuspended in mitochondria isolation buffer without BSA and pelleted at 10 000 g for 10 min.

### Proteinase K accessibility assay

Mitochondria were isolated as described in ^40^ and the proteinase K accessibility assay was performed as previously described ^41^. Briefly, after 24 h transient overexpression of NERCLIN cells from at least two 15-cm dishes were harvested. Cells were washed once with cold PBS and spun at 500 g for 3 min at 4°C. The cell pellet was resuspended in 10 mL of ice-cold extraction buffer (10 mM Tris·Mops, pH 7.4, 1 mM EGTA·Tris, pH 7.4, 0.2 M sucrose, pH adjusted to 7.4) with 1x protease inhibitor cocktail (Thermo Scientific), and cells were disrupted using a glass-glass homogenizer. Unbroken cells were spun by centrifugation at 600 g for 10 min at 4°C, and the pellet was subjected to the second round of homogenization. Mitochondria were recovered from the supernatant of both rounds of homogenization by centrifugation at 7000g for 10 min at 4°C, and the pellet was washed once with a cold extraction buffer and centrifuged again. The final pellet was resuspended in 200 μL of extraction buffer and the mitochondrial protein concentration was determined with the Pierce BCA Protein Assay Kit (Thermo Scientific). For the Proteinase K accessibility assay, 50 μg of mitochondria were resuspended in 100 μL of extraction buffer (untreated), extraction buffer and Proteinase K (protease-treated mitochondria), 2 mM Hepes, pH 7.4 and Proteinase K (protease-treated mitoplasts), or 2 mM Hepes, pH 7.4, 0.1% Triton X-100 and Proteinase K (protease- and detergent-treated mitoplasts). 100 μg/mL Proteinase K (Thermo Scientific) was used in all cases. Samples were incubated on ice for 30 min and the reaction was inactivated with a final concentration of 1mM PMSF. Samples were precipitated with 10% trichloroacetic acid, washed once with cold acetone and resuspended in 50 μL of 4 × Laemmli sample buffer containing 4% (vol/vol) β-mercaptoethanol and 1x HALT. Subsequently, 10 μL of each sample were boiled at 95°C for 5 minutes and separated by SDS/PAGE, followed by Western blotting analysis.

### Prediction of NERCLIN structure

NERCLIN protein secondary structure was predicted using Chou and Fasman Secondary Structure Prediction server, CFSSP (https://www.biogem.org/tool/chou-fasman), Jpred4 (http://www.compbio.dundee.ac.uk/jpred4)^42^, secondary structure prediction method GOR4^43^, PSIPRED server^44^ and PHD (https://npsa-prabi.ibcp.fr/cgi-bin/npsa_automat.pl?page=/NPSA/npsa_phd.html). Tertiary structure of human NERCLIN was predicted by SSpro server (http://scratch.proteomics.ics.uci.edu/). For the analysis mitochondrial targeting sequence (1-32 amino acids) was removed from the protein sequence.

### DNA and RNA analysis

For expression analysis human multiple tissue cDNA panel (MTC™ Clontech, 636742) and human normal brain tissue qPCR array (OriGene Technologies, HBRT101) were used. 1-3 ng of cDNA was used for real-time PCR assay. After completion of PCR amplification of the samples from a human multiple tissue cDNA panel, the PCR products were loaded on 0.7% agarose gel for electrophoresis.

### Real-time PCR assay

Total RNA was extracted from cultured cells using Mini spin kit (Macherey-Nagel). RNA extracted from canine fibroblasts was obtained from Prof. Hannes Lohi group. 1000 ng of RNA was reverse transcribed using Maxima First Strand cDNA Synthesis Kit for RT-qPCR (Thermo Fisher scientific). Real-time PCR analysis was done by DyNAmo Flash SYBR Green qPCR Kit (Thermo Fisher scientific) using CFX96™ Real-Time PCR Detection System (Bio-Rad). The PCR program started with 95°C for 7 min followed by 40 cycles at 95°C for 10 s and 60°C for 30 s. Beta-2-Microglobulin (B2M) was used as a reference gene for normalization and mRNA expression level was calculated using the comparative Ct (threshold cycle) method. All primer sequences are in the Table 1.

**Table 1.**
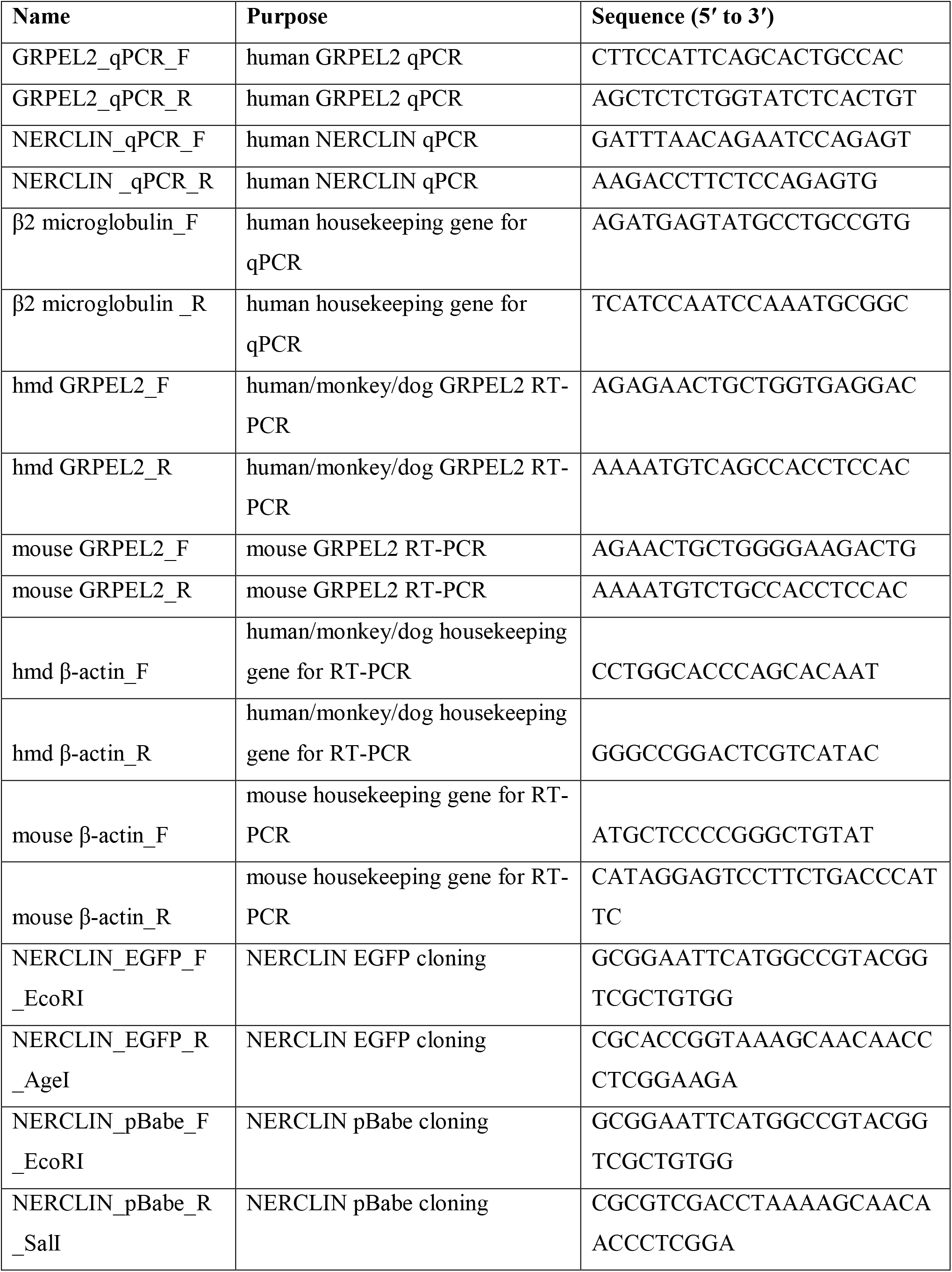
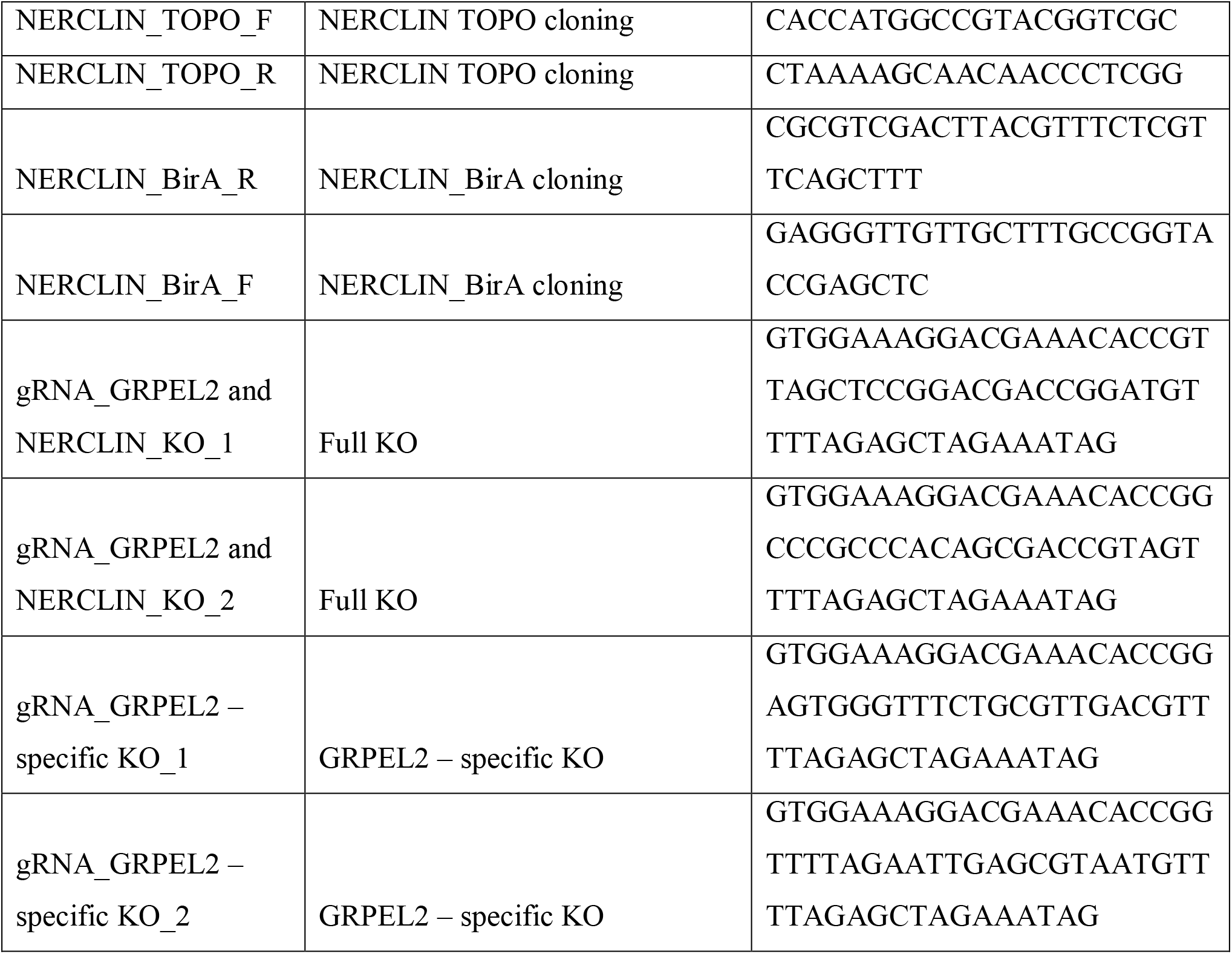
List of primers used in this study

### Mass spectrometric analysis of lipids

HEK cells were grown on 15 cm dishes. Lipids were extracted from the cell pellets or isolated mitochondria according to the Folch method^45^. The samples were analyzed with electrospray ionization-tandem mass spectrometry (ESI-MS/MS) using 6410 Triple Quadrupole LC/MS (Agilent Technologies, CA, USA). The final lipid extracts in chloroform:methanol (1:2, v:v) were spiked with internal phospholipid standards or Ceramide/Sphingoid Internal Standard Mixture I (Avanti Polar Lipids, USA) and 1% NH4OH (Merck) right before infusion into the MS at a flow rate of 10 μL/min. The internal phospholipid standards were phosphatidylcholine (PC) 14:1/14:1, PC 20:1/20:1, PC 22:1/22:1; phosphatidylethanolamine (PE) 14:1/14:1 and PE 20:1/20:1; phosphatidylserine (PS) 14:1/14:1; phosphatidylinositol (PI) 16:1/16:1; and sphingomyelin (SM) 18:1/17:0 (from Avanti Polar Lipids or prepared in house). The MS source temperature was 250°C and instrument nitrogen was used as the nebulizing (40 psi) and the drying gas (3 L/min).

Precursor ion scans were employed to produce lipid class specific scans as follows: *m/z* 184 for PC and SM; *m/z* 241 for PI; *m/z* 264 for ceramide (Cer) and hexosylceramide (HexCer); and *m/z* 369 for cholesterol ester (CE) as ammonium adducts^46–48^. Neutral loss scans were employed for PE (loss of 141 amu) and PS (loss of 87 amu)^46^. PE plasmalogens (PEp) were identified with fragment-specific scans for the vinyl ether chains at the *sn*-1 position (e.g., *m/z* 364, 390, and 392 for 16:0p, 18:1p, and 18:0p, respectively) and analyzed from the MS-scan^49^. Triacylglycerol (TAG) species were analyzed as ammonium adducts from the MS+ scan^50^. For CL and PG species analysis, 400 μL of the lipid extract was spiked with CL 14:0/14:0/14:0/14:0 and PG 20:1/20:1 (Avanti Polar Lipids) internal standards, methylated^51^ and the lipids were extracted as above^45^. The samples were then run on ACQUITY Ultra Performance LC system coupled to ESI source of Quattro Micro triple quadrupole MS (Waters, Manchester, UK) as described previously^52^ and the spectra were extracted from the chromatogram. The CL were detected by MS+ scanning, and the PG species were selectively detected by scanning for the neutral loss of 203 amu^53^ All lipid species were identified and quantified using Lipid Mass Spectrum Analysis software^54^ and internal and external standards. The lipid species are marked as follows: [lipid class] [sum of acyl chain carbons]:[sum of acyl chain double bonds] (e.g., PC 34:1). The data are described as molar percentages (mol%) for each individual species relative to its lipid class.

The lipidomics data were analyzed with R (https://www.R-project.org/) using limma^55^. The raw intensities were normalized with the voom-function. The analysis was done using linear regression (lmFit), with empirical Bayesian statistics (eBayes).

### Fluorometric measurement of total CL

To measure total CL in cells and isolated mitochondria fluorometric CL assay kit (BioVision, K944) was used. For sample preparation, cells or mitochondrial pellets were resuspended in the CL assay buffer and lysed by sonication. The debris was removed by centrifugation at 10 000 rpm for 10 min at +4°C. Then, the CL measurement was performed according to the manufacturer’s instruction.

### Immunocytochemistry

143B cells were transiently transfected with pDsRed2-Mito Vector (Clontech, 632421) to label mitochondria. In the experiments for studying intracellular localization of NERCLIN, cells were co-transfected with the NERCLIN-EGFP construct. In the experiments with BirA* fused proteins, cells were co-transfected with NERCLIN-BirA*. After 24 h of transfection cells were fixed with 4% paraformaldehyde for 10 min at RT and washed with PBS. Then cells were permeabilized with Triton X-100 for 15 min at RT, washed and blocked with 5 % BSA for 2 h at RT. Cells were then incubated with corresponding primary antibody against GRPEL2 (Novus Biological, NBP1-85099) in a blocking buffer overnight at +4 °C. After washing cells were incubated with secondary antibody for 1 h at RT (Alexa Fluor 488 goat anti-rabbit, Invitrogen, R37116). Finally, cells were washed, mounted and imaged with Axio Observer Z1 (Zeiss).

### Plasmid construction

To clone the NERCLIN cDNA we extracted total RNA from 143B cells using Mini spin kit (Macherey-Nagel). Then RNA was reverse transcribed using Maxima First Strand cDNA Synthesis Kit for RT-qPCR (Thermo Fisher scientific). The cDNA was PCR amplified using specific primers for TOPO cloning. Following gel electrophoresis the correct band was cut and the DNA was extracted from the gel for TOPO cloning. The TOPO construct containing NERCLIN was used as a template for cloning NERCLIN into EGFP or pBabe vectors.

To fuse BirA* to C-terminus of NERCLIN we used PCR overlap extension. pHA-BirA* and pBabe-NERCLIN constructs were used the templates. The resulted fused construct was inserted into pBabe vector using *EcoR*I and *Sal*I restriction sites. GFP-BirA* and AIF-BirA* plasmids were as in ^23^.

### Generation of gene knockouts

CRISPR/Cas9 was used to generate knockout HEK293 cells. Cells were co-transfected with two gRNA transcriptional cassettes prepared by PCR and CAG-Cas9-T2A-EGFP plasmid (Addgene #7831) as described elsewhere^56^. To generate GRPEL2 specific knockout cell line one of the guide RNAs was targeted to the intron 2 region and the other to intron 3 of the *GRPEL2* gene. After 24 h of transfection GFP-positive cells were sorted by FACS and single cell clones generated. The sequences of gRNAs are in the Table 1.

### Western blotting

Whole cell or mitochondrial samples were lysed in a RIPA buffer supplemented with protease inhibitor cocktail (Thermo Scientific). Following 10 min incubation on ice, the samples were centrifuged at 14 000 g for 10 minutes at +4°C. Protein concentration was measured by bicinchoninic acid (BCA) assay (Thermo Fisher). Protein lysates were supplemented with a laemmli sample buffer (Bio-Rad) and resolved on 10 % Mini-PROTEAN TGX Precast Gels (Bio-Rad). To detect OPA1 isoforms proteins were resolved on 7.5% Mini-PROTEAN TGX Precast Gels (Bio-Rad). Then proteins were transferred to a 0.2-μM PVDF membrane by Trans-Blot Turbo Transfer System (Bio-Rad). The membranes were blocked in 5% milk in TBS–Tween 20 (0.1%) for 1 h at room temperature. Proteins were immunoblotted with the indicated primary antibodies in TBS–Tween 20 (0.1%) containing 1% BSA for overnight, at 4 °C. Following primary antibodies were used: anti-GRPEL2, 1:1000 (Novus, 90536); anti-TFAM, 1:1000 (Abcam 131607) anti-PHB, 1:1000 (Boster Bio PA1932); anti-PHB2, 1:1000 (BioLegend 611802); anti-beta-tubulin, 1:1000 (Cell Signaling 2146); anti-NDUFA9 1:2000 (Abcam 14713); anti-SDHA 1:2000 (Abcam 14715); anti-COX1 1:2000 (Abcam 14705); anti-HSP60 1:2000 (Santa Cruz 1052); anti-LC3B 1:1000 (Novus biologicals 600-1384); anti-p62 1:5000 (Abnova 2C11); anti-ATP5A 1:1000 (Abcam 14748); CHCHD10 1:1000 (Sigma HPA003440); anti-COX2 1: 1000 (GeneTex GTX62145); anti-TOM40 1:1000 (Santa Cruz 11414); anti-OPA1 1:1000 (BD Biosciences 612606). Membranes were extensively washed and probed with HRP conjugated secondary antibodies against mouse, rabbit or goat IgG (Jackson ImmunoResearch) at 1:5000 in TBS–Tween 20 (0.1%) containing 1% BSA and washed again. Chemiluminescence was captured by Chemidoc imaging system (Bio-Rad). Quantification of the bands was performed by Image Lab Software (Bio-Rad).

### BioID analysis

143B cells were grown on 15 cm dishes. The cells were transfected with corresponding BirA* fusion constructs using jetPRIME according to the manufacturer’s manual. Following 24 h of transfection, 50 μM biotin was added to each plate and allowed to biotinylate the proximal proteins for the next 24 h. The plates were washed with PBS, and cells scraped, pelleted and frozen at −80°C. For each sample, four biological replicates were analyzed. Biotinylated proteins were extracted using streptavidin beads and analyzed by mass spectrometry as previously described^20^. The mass spec analysis was done in technical duplicates. The mass spectrometry data were analyzed as previously described ^20, 24^.

### Transmission electron microscopy

HEK293 cells were cultured on glass coverslips in 6-well plates for 24 h, then transfected with the corresponding construct. After 48 h of transfection the cells were fixed with a solution of 2% glutaraldehyde in 0.1 M sodium cacodylate buffer at pH 7.4 for 25 minutes at room temperature. Then the cells were washed two times for 3 min with 0.1 M sodium cacodylate buffer (pH 7.4). Fixed cells were then processed according to the standard protocol at Electron Microscopy Unit of the Institute of Biotechnology, University of Helsinki. Images were acquired with the Jeol JEM-1400 transmission electron microscope. ImageJ software was used to quantify mitochondrial major axis length (mitochondrial length) or cristae width.

### Statistical Analysis

All data are presented as mean ± standard deviation (SD). For the statistical analyses with two samples, student’s unpaired two-tailed t-tests or one-way ANOVA analysis were performed using Graph Pad Prism Software. All the graphs were prepared with Graph Pad Prism Software.

## ACKNOWLEDGEMENTS

Riitta Lehtinen, Sanna Sihvo and Tarja Grundström are thanked for excellent technical assistance. Electron Microscopy Unit at the Institute of Biotechnology. Docent Marjo Hytönen and Prof. Hannes Lohi are thanked for sharing the canine RNA sample. Group of Thomas McWilliams is thanked for sharing anti-OPA1 antibody. This work was supported by European Research Council (Grant no. 637458), Academy of Finland, University of Helsinki and Sigrid Juselius Foundation.

## AUTHOR CONTRIBUTIONS

SK conceptualized and supervised the project, designed and performed experiments and analyzed data. SB, RTM, PM, JR, YY performed experiments. XL, MV performed BioID mass spec analysis. PS, MH and RK performed lipid analysis. JK statistically analyzed lipidomics data. SK and HT wrote the manuscript. HT supervised the study and provided funding and resources. All authors commented on the manuscript.

## CONFLICT OF INTEREST STATEMENT

The authors declare that they have no conflict of interest.

## SUPPLEMENTARY FIGURE LEGENDS

**Supplementary Figure 1:**
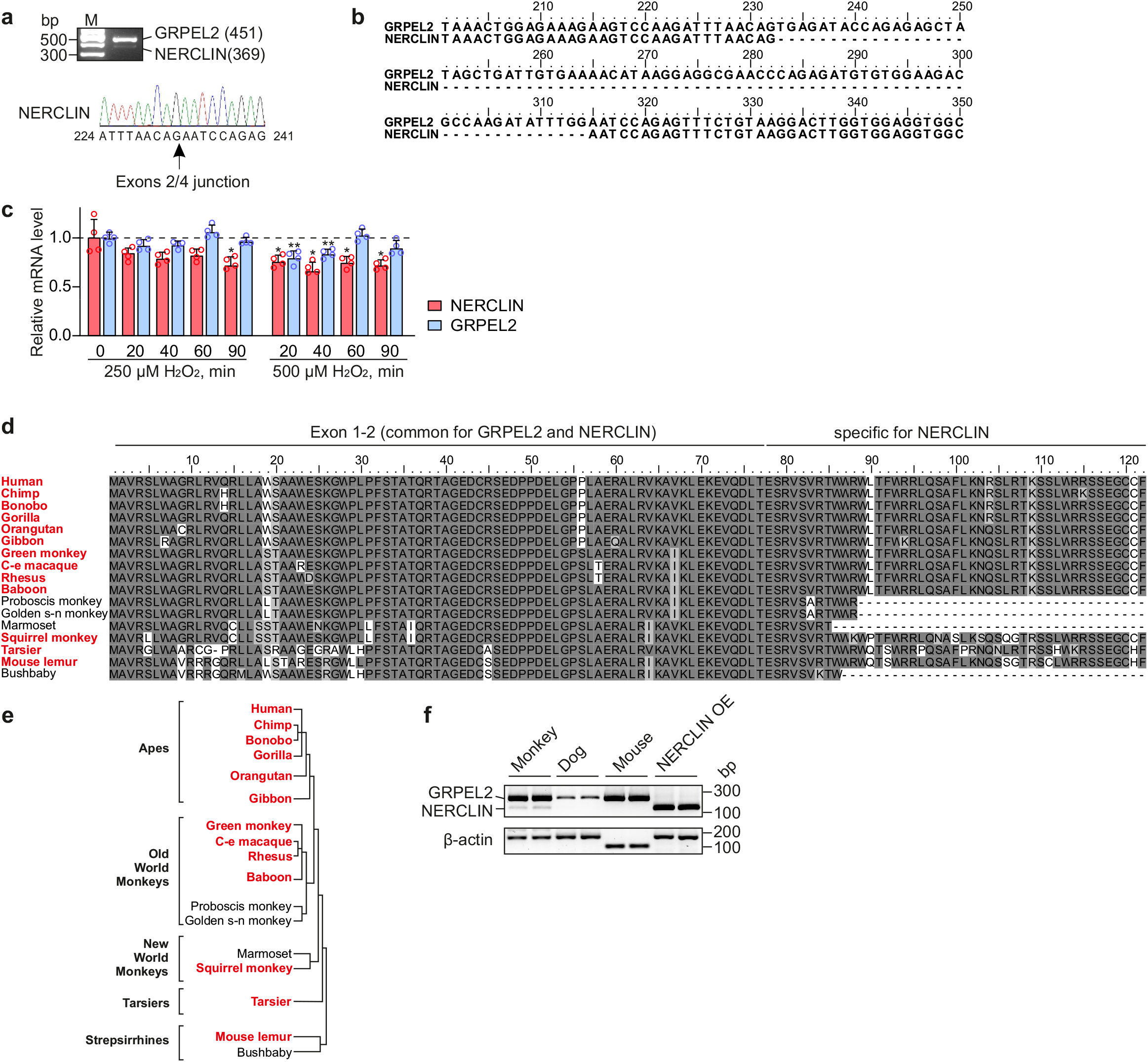
Conservations of NERCLIN protein in primates and other species. **a** RT-PCR co-amplification of two GRPEL2 transcripts in 143B cells. Primers spanning NERCLIN cDNA were used (upper panel). The expected band sizes in bp are indicated in brackets. M, DNA ladder. RT-PCR products were cloned and sequenced. DNA sequencing chromatogram indicating junction of exon 2 and 4 in GRPEL2 variant (lower panel). **b** cDNA sequence alignment of GRPEL2 and NERCLIN lacking exon 3 (233–724 bp). The RT-PCR products as in **(a)** were used for nucleotide sequencing. **c** mRNA levels of NERCLIN or GRPEL2 in HEK293 cells exposed to H_2_O_2_ as determined by qPCR assay (n=4). Data are presented as mean ± SD. *P < 0.05, **P < 0.01, as compared to untreated cells (unpaired t-tests). **d** Multiple sequence alignment of GRPEL2 protein transcript skipping exon 3 in primates. Identical residues are highlighted in dark gray, similar residues are in light gray. Species that are predicted to have NERCLIN are in red. C-e macaque, crab-eating macaque; Golden s-n monkey, golden snub-nosed monkey. **e** Phylogeny tree of primates analyzed in **(d)**. Species that are predicted to have NERCLIN are in red. **f** NERCLIN transcript detection in monkey kidney cells (COS7), dog fibroblasts and mouse liver. RT-PCR products of GRPEL2 and NERCLIN, 260 bp and 168 bp respectively were detected on the gel. HEK293 cells overexpressing NERCLIN were used as a positive control. B-actin was used as a loading control.

**Supplementary Figure 2:**
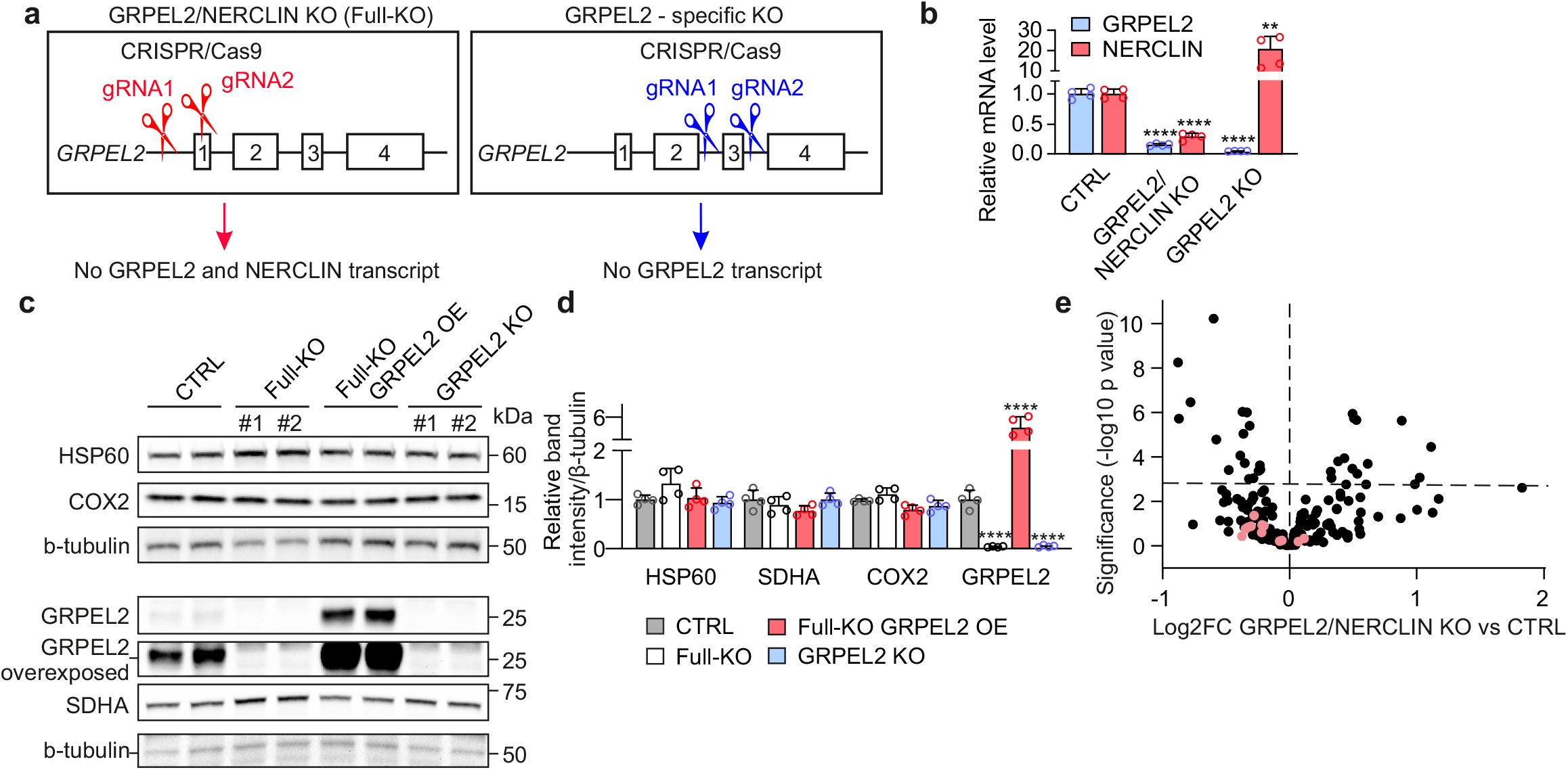
GRPEL2/NERCLIN knockout or GRPEL2 knockout does not cause mitochondrial abnormalities or cardiolipin depletion. **a** CRISPR/Cas9 approach using guide RNAs (gRNAs) targeted to specific regions of the *GRPEL2* gene to generate GRPEL2/NERCLIN knockout or GRPEL2 knockout HEK293 cells. **b** GRPEL2 or NERCLIN transcript levels in GRPEL2/NERCLIN knockout or GRPEL2 knockout cells determined by qPCR assay (n=4). **c** Western blot analysis of mitochondrial proteins in GRPEL2/NERCLIN knockout (Full-KO) or GRPEL2 knockout cells. **d** Quantification of western blot images presented in **(c)** (n=4). **e** Volcano plot compares the lipid species composition of GRPEL2/NERCLIN knockout (KO) HEK293 cells and control cells (CTRL). The dashed line indicates a false discovery rate (FDR)-corrected P-value of 0.05. CL species are in light red. The data are presented as mean ± SD. **P <0.01, ***P <0.001, ****P < 0.0001, ns, not significant as compared to the control cells (unpaired t-tests).

**Supplementary Figure 3:**
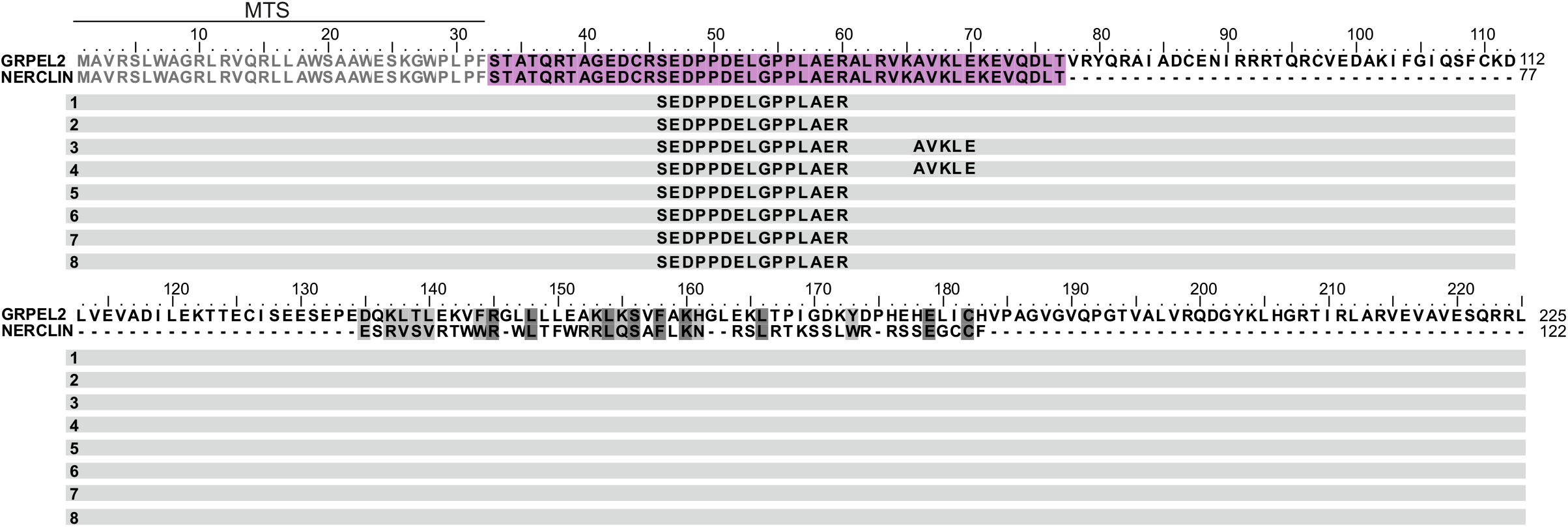
NERCLIN does not show proximity to GRPEL2. Peptide spectrum matches of GRPEL2 in 143B cells overexpressing NERCLIN-BirA* identified by mass spectrometry in BioID analysis. Amino acid sequence alignment of human GRPEL2 and NERCLIN. Identical residues are highlighted in dark gray, similar residues are in light gray. Amino acid sequence common for GRPEL2 and NERCLIN is in violet. Mitochondrial targeting sequence, MTS (1-32 amino acids) is in gray. Each line represents replicate for mass spectrometry.

**Supplementary Figure 4:**
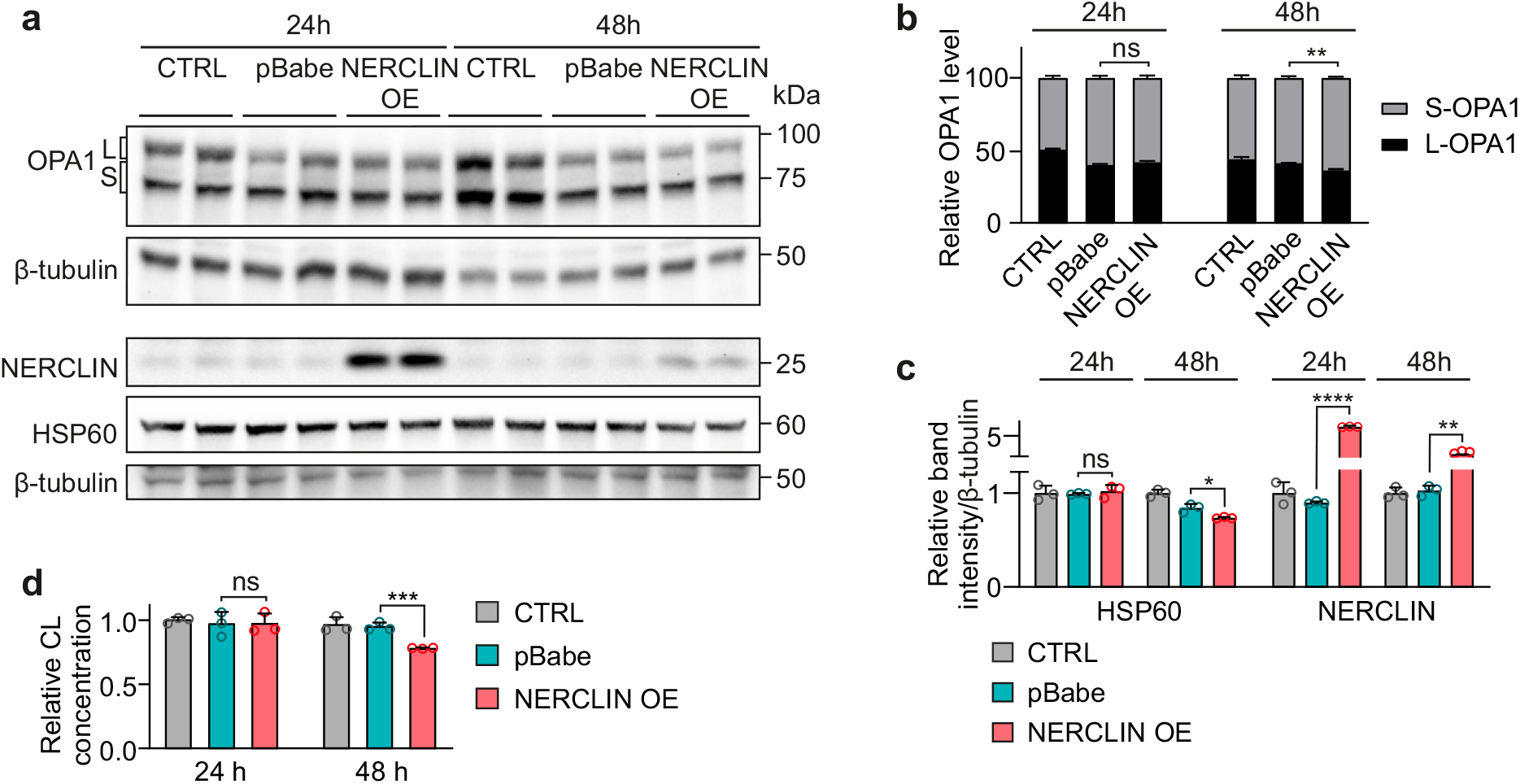
Short overexpression of NERCLIN does not cause mitochondrial fragmentation or changes in CL content. **a** Western blot analysis of HEK293 cells transfected with NERCLIN plasmid (NERCLIN OE) or with empty vector (pBabe) for 24 h or 48 h. **b** Relative levels of short and long OPA1 isoforms determined by western blot analysis in **(a)**. L, long OPA1 isoform, S, short OPA1 isoform (n=3). **d** Quantification of western blot images presented in **(a)**. Protein expression levels are normalized to β-tubulin level (n=3). **c** Total CL concentration in HEK293 cells transfected with NERCLIN or empty vector (pBabe) as determined by fluorometric assay (n=3). In all graphs data are presented as mean ± SD. *P < 0.05, **P < 0.01, ***P < 0.001, ****P < 0.0001, ns, not significant as compared to the cells transfected with empty vector (unpaired t-tests).

**Supplementary Figure 5:**
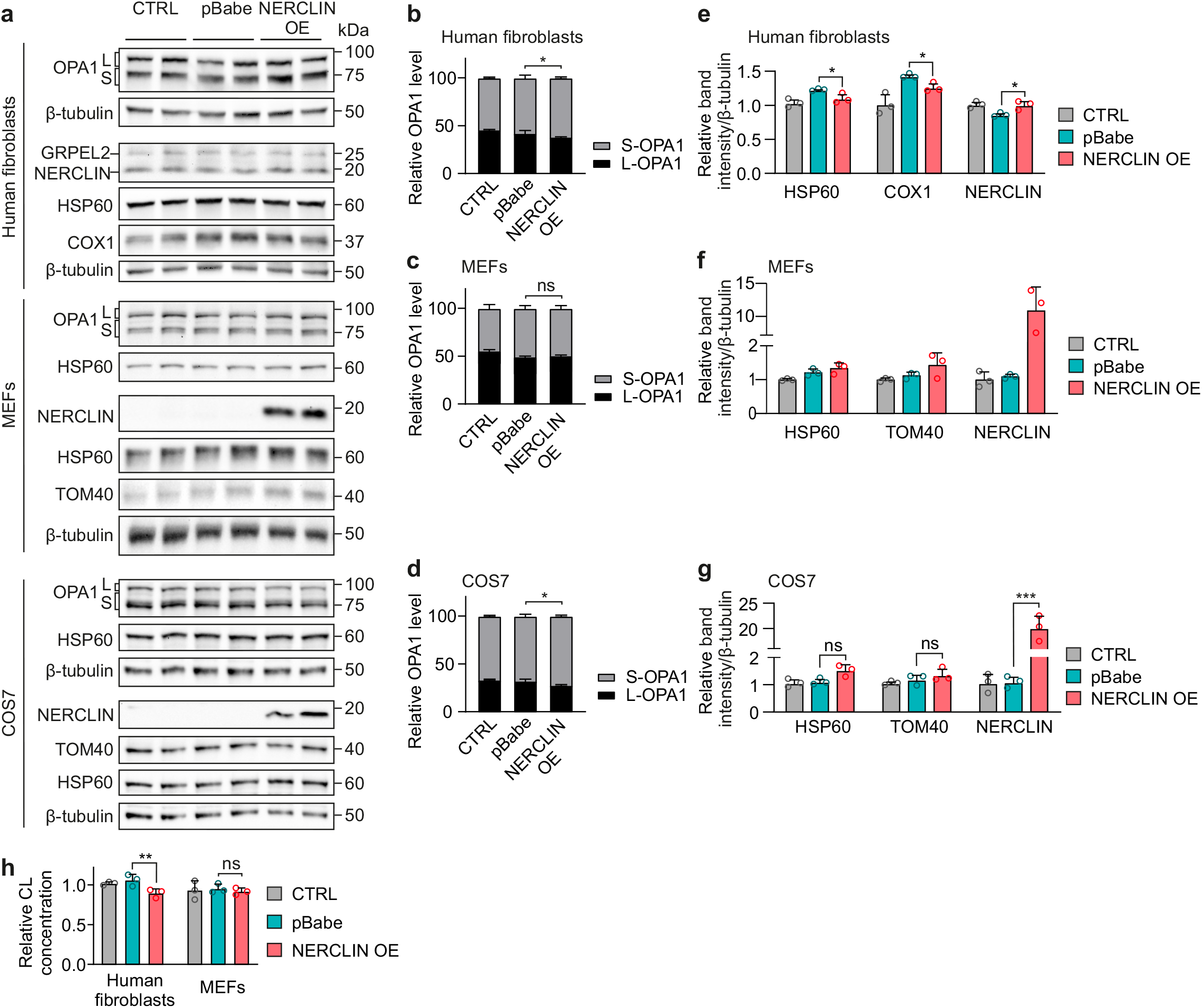
Overexpression of NERCLIN specifically affects mitochondria in primate cells. **a** Western blot analysis of human fibroblasts, mouse embryonic fibroblasts (MEFs) or African green monkey kidney fibroblasts (COS7) overexpressing NERCLIN. Cells were transiently transfected with NERCLIN plasmid (NERCLIN OE) or with an empty vector (pBabe) for 48 h. CTRL, non-transfected cells. **b-d** Relative levels of short and long OPA1 isoforms determined by western blot analysis in **(a)**. L, long OPA1 isoform, S, short OPA1 isoform (n=3). **e-g** Quantification of western blot images presented in **(a)**. Protein expression levels are normalized to β-tubulin level (n=3). **h** Total CL concentration in HEK293 cells transfected with NERCLIN or empty vector (pBabe) as determined by fluorometric assay (n=3). In all graphs data are presented as mean ± SD. *P < 0.05, **P < 0.01, ***P < 0.001, ns, not significant as compared to the cells transfected with empty vector (unpaired t-tests).

## SUPPLEMENTARY TABLES

**Supplementary Table 1. Identified unique interactors of NERCLIN**

**Supplementary Table 2. Lipidomics analysis of cells overexpressing NERCLIN**

## Notes

### Competing Interest Statement

The authors have declared no competing interest.

